# Sensitizing solid tumors to CAR-mediated cytotoxicity using synthetic antigens

**DOI:** 10.1101/2021.12.11.472238

**Authors:** Lena Gamboa, Ali H. Zamat, Daryll Vanover, Chloé A. Thiveaud, Hannah E. Peck, Hathaichanok Phuengkham, Anirudh Sivakumar, Adrian M. Harris, Shreyas N. Dahotre, Fang-Yi Su, Philip J. Santangelo, Gabriel A. Kwong

## Abstract

CAR T cell immunotherapy relies on CAR targeting of tumor-associated antigens, yet heterogenous antigen expression, interpatient variation, and off-tumor expression by healthy cells remain barriers. Here, we develop synthetic antigens to sensitize solid tumors for recognition and elimination by CAR T cells. Unlike tumor-associated antigens, we design synthetic antigens that are orthogonal to endogenous proteins to eliminate off-tumor targeting and that have a small genetic footprint to facilitate efficient tumor delivery to tumors by viral vectors. Using the RSV-F camelid single-domain antibody (VHH) as a synthetic antigen, we show that adoptive transfer of αVHH CAR T cells to mice bearing VHH expressing tumors reduced tumor burden in multiple syngeneic mouse models of cancer, improved survival, induced epitope spread, and protected against tumor rechallenge. Our work supports *in situ* delivery of synthetic antigens to treat antigen low or negative tumors with CAR T cells.

## MAIN TEXT

Despite the therapeutic efficacy of chimeric antigen receptor (CAR) T cells in treating certain B cell malignancies (*1*), their clinical success against solid tumors remains limited (*2–4*). Barriers to effective CAR T cell therapy for solid tumors are multifaceted and include physical barriers that limit T cell infiltration, the presence of an immunosuppressive microenvironment that promotes T cell dysfunction, and the difficulty in identifying antigens that elicit therapeutic responses upon targeting (*5*). Unlike liquid tumors like acute lymphoblastic leukemia (ALL) where most malignant cells express the B cell marker CD19, antigen selection remains a bottleneck to achieving effective CAR T cell therapy for solid tumors. Heterogenous antigen expression within a tumor and interpatient variation in expression limit the broad applicability of a single CAR construct. Human breast cancers, for example, are classified as estrogen receptor (ER)-positive when as few as 1% of tumor cells express ER (*6*). Additionally, low antigen density and antigen loss, such as through trogocytosis and acquired mutations, render CAR T cells ineffective over time and raise concerns with antigen escape (*7–9*). Thus, genome wide activation of endogenous genes using CRISPRa has been proposed to augment tumor antigen expression (*10*). Moreover, emerging strategies to mitigate antigen escape include the use of universal targeting motifs to redirect CAR T cells against multiple antigens using adaptor molecules (*11–14*), or the use of CAR T cells engineered with constitutive or heat triggered circuits that enable spatial control of bispecific T cell engagers that target NKG2D ligands to redirect T cells against tumor cells that do not express the CAR antigen (*15, 16*).

The problem of antigen heterogeneity is compounded by off-tumor toxicities arising from shared expression of target antigens with healthy tissue (*17–19*). CD19 CAR T cell therapy, for example, leads to the depletion of healthy B cells, resulting in hypogammaglobulinemia and cytopenia that typically requires lifelong IgG replacement therapy while also carrying the risk of neurotoxicity due to low-level expression of CD19 on healthy brain mural cells (*20*). T cells targeting other endogenous tumor-associated antigens (TAAs) (e.g., HER2, CAIX) show cross-reactivity with healthy cells in vital organs (*4, 21*), leading to on-target, off-tumor toxicities that limit safe and effective clinical translation (*22*). To address toxicity concerns with the overlap in antigen expression between malignant and healthy tissue, numerous approaches are being evaluated, including the use of thermally controlled circuits to focus CAR T cell cytotoxic activity to the tumor site (*15, 23*) and the integration of combinatorial antigen sensing circuits to enhance the ability of engineered T cells to discriminate malignant cells from normal tissue expressing a single epitope (*14, 24*). In addition to challenges in identifying solid tumor antigens with therapeutic potential, strategies that increase the therapeutic index of CARs by optimizing CAR affinity (*25, 26*), as well as reports demonstrating the effects of linker length and intracellular domains on CAR function (*27, 28*), highlight the need to carefully optimize CAR constructs for each potential antigen target. This iterative process is costly and time-consuming, limiting downstream clinical implementation of CAR T cells for a broad range of tumor types. To overcome challenges in identifying TAAs and alleviate the need to validate CARs for each target antigen candidate, direct delivery of CD19 via oncolytic virus or GFP by tumor-colonizing bacteria enabled tumor recognition by a single CAR T cell construct (*29–31*). Collectively, these efforts underscore the need to develop strategies that reduce the bottleneck of antigen selection and enable CAR T cell recognition of tumors otherwise lacking targetable antigens.

We developed synthetic antigens to sensitize solid tumors to CAR T cell mediated immunity agnostic to a tumor’s endogenous antigen expression profile. Ideally, a synthetic antigen would be characterized by several key features: it should (1) be orthogonal to endogenous proteins to minimize off-tumor toxicity, (2) be genetically encoded for delivery by viral and nonviral approaches, and (3) be small and compact to facilitate gene transfer. Additionally, it should (4) be stably expressed to facilitate recognition and (5) be targetable by single-chain variable fragments (scFvs) to enable engineering of cognate CAR T cells against them. Considering these criteria, we repurposed a xenogeneic protein, the antigen binding fragment of the heavy-chain-only camelid antibodies (VHH), as a synthetic antigen. Its small genetic footprint (~375 bp), thermal stability, protease resistance, and stability at extreme pH have facilitated numerous *in vivo* applications (*32–34*), including as neutralizing agents (*35, 36*), fusion proteins for imaging (*37, 38*), payload carriers for targeted vaccine delivery (*39, 40*), and chimeric antigen receptors in lieu of the traditional scFv domain (*41*). We demonstrate stable expression of VHH on the surface of various tumor types, including breast, colon, and lung cancer, following mRNA or AAV delivery, which subsequently led to recognition and killing by αVHH CAR T cells. Moreover, we show αVHH CARs are well tolerated in naïve C57BL/6J mice at the doses tested by serum chemistry. Adoptive transfer of αVHH CAR T cells to mice bearing VHH-treated tumors reduced tumor burden in multiple syngeneic models of cancer, improved survival, induced epitope spread and prevented tumor growth after rechallenge. Our data shows that sensitizing tumors to CAR-mediated cytotoxicity using synthetic antigens could potentiate antitumor immunity and improve responses against solid tumors.

## RESULTS

### Synthetic antigens are expressed by multiple tumor types following mRNA delivery in vitro

To achieve surface expression of VHH for downstream CAR targeting, we designed and tested several synthetic antigen sequences comprised of a membrane anchor, a linker, and a recognition domain (**Fig. 1a**). We first compared the CD4 transmembrane (CD4TM) domain (*42*) to the glycosylphosphatidylinositol (GPI) anchor from the decay accelerating factor (DAF) (*42–44*) as membrane anchors for RSV-F VHH. In wildtype E0771, MC38, and A549 cancer cells, we quantified surface VHH expression by surface staining and flow cytometry following mRNA transfection (**Fig. 1b**). Maximum surface expression, as measured by median fluorescence intensity (MFI), was ~10-fold higher for GPI-anchored VHH compared to CD4TM-anchored VHH across three tumor cell lines (**Fig. 1c**). Moreover, VHH expression was maintained above background for 5–12 days for the GPI-anchored construct, whereas CD4TM-VHH remained detectable above background for 2–4 days (**Fig. 1c**). Due to the higher and more sustained overall expression of the GPI-anchored constructs, we chose to utilize GPI as the membrane anchor for synthetic antigens.

**Fig. 1.**
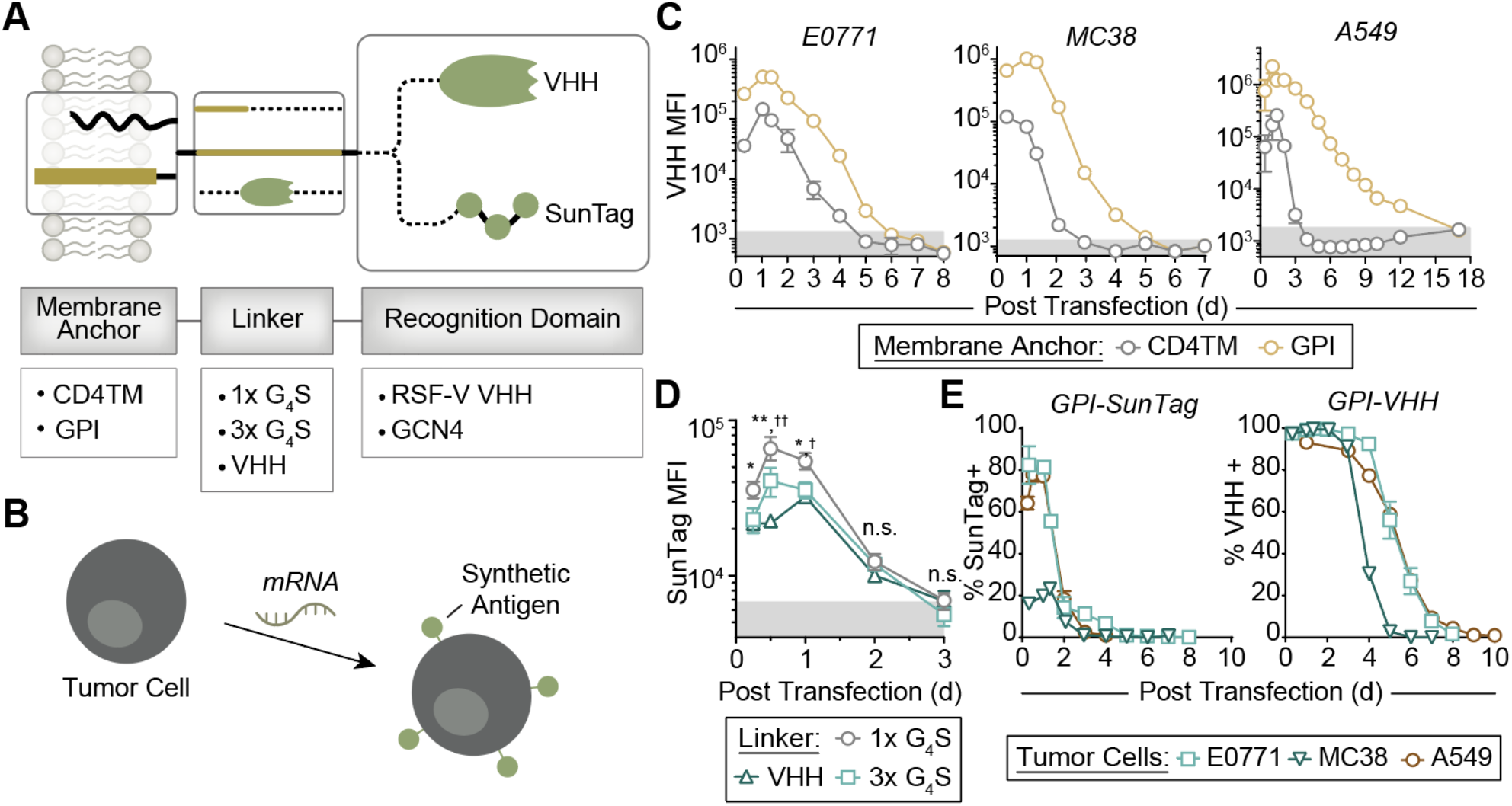
Expression kinetics of synthetic antigen constructs. (a) mRNA constructs tested consist of a membrane anchor, a linker, and a recognition domain. These constructs are (b) transfected into tumor cells to achieve synthetic antigen expression. (c) Expression kinetics of GPI-anchored and CD4TM-anchored synthetic antigens consisting of a 1x G_4_S linker and the RSV-F VHH recognition domain on the surface of indicated tumor cells. (d) Expression kinetics of mRNA constructs with a GPI-anchor, SunTag recognition domain, and indicated linker domains. Two-way ANOVA, mean ± s.d. is depicted; n = 4 (for 1x G_4_S vs 3x G_4_S: \**p* < 0.05, \*\**p* < 0.01; for 1x G_4_S vs 3x G_4_S: † *p*< 0.05, †† *p* < 0.01). (e) Expression kinetics of GPI-anchored SunTag or VHH on the surface of indicated tumor cell lines.

Next, to evaluate the stability of the VHH recognition domain relative to a recognition domain with a smaller genetic footprint, we compared expression kinetics of the VHH protein (375 bp) to a short repeating peptide array, SunTag (201 bp). SunTag is a repeat of the 19-aa peptide epitope EELLSKNYHLENEVARLKK of the yeast transcriptional factor GCN4 originally integrated into a protein tagging system due to its small genetic footprint, orthogonality to naturally occurring sequences in the genome, and ability to bind to its cognate scFv with high affinity and specificity (*45*). Despite their shorter sequences, all three SunTag constructs tested, each linked to the GPI anchor by either a 1x repeat of the flexible G_4_S peptide, a 3x repeat of G_4_S, or a protein (i.e., VHH) linker, became indistinguishable from background fluorescence by 3 days post transfection in A549 cells (**Fig. 1d**). Moreover, while VHH-linked SunTag became undetectable by day 3 (**Fig. 1d**, triangles), only the VHH linker (and not its tethered SunTag) remained detectable on 96.7% of cells by flow cytometry 5 days post transfection (**Fig. S1**), demonstrating that VHH is more stable than SunTag when expressed on the surface of tumor cells. By comparison, the MFI of GPI-anchored VHH was detectable on A549s for at least 12 days (**Fig. 1c**). Half-maximal expression for GPI-anchored, 1xG_4_S-linked SunTag was reached on average 1.8 days post transfection across E0771, MC38, and A549 tumor lines, while VHH reached half-maximal expression on average by 4.8 d (**Fig. 1d**). Altogether, GPI-anchored VHH showed sustained surface expression on multiple cell lines, both murine and human, for at least 5 days (**Fig. 1e, Fig. S1-S2**). For comparison, the Kb-SIINFEKL pMHC complex has a half-life of ~8 hrs on the surface of MC38 cells (**Fig. S3**), while various human immune cells such as DCs, B cells, and monocytes have a reported half-life of the HLA-A*02:01-gp100_154-162_ complex of 1.5 –22.5 hrs (*46*), demonstrating that VHH can be expressed for biologically relevant durations following transient mRNA transfection.

### αSunTag and αVHH CARs recognize and kill tumor cells expressing their cognate synthetic antigen in vitro

To target synthetic antigen expressing tumor cells, we designed murine CAR T cells by fusing the anti-VHH and anti-GCN4 scFv sequences to the murine CD8α hinge and transmembrane domain, CD28 costimulatory domain, and the CD3ζ signaling domains (**Fig. 2a-b**). Similarly, human CARs comprised of either an anti-VHH or anti-GCN4 scFv, followed by the human CD8α hinge and transmembrane domain, 4-1BB costimulatory domain, and CD3ζ cytolytic domain (**Fig. S4a**). Both VHH and SunTag CARs were stably expressed on the surface of both primary murine and primary human T cells (**Fig. 2c, Fig. S4b**) following viral transduction. To test their antigen-dependent activity, murine CAR T cells were co-incubated with VHH- or SunTag-transfected E0771 cells (**Fig. 2d**). Flow cytometry analysis of CD25 and CD69 activation markers revealed that greater than 94% of CAR T cells expressed both activation markers following exposure their cognate synthetic antigen (**Fig. 2e**). Moreover, following a 24 hr co-culture at a 2:1 effector:target ratio, αVHH and αSunTag CAR T cells secreted interferon gamma (IFN-γ) at significantly higher levels over wildtype T cells when co-cultured with VHH- or SunTag-expressing tumor cells (\*\*\*\**p* < 0.001), respectively, but no significant elevation was detected when cultured alone or with tumor cells expressing a mismatched synthetic antigen (**Fig. 2f**). Similarly, we observed significantly elevated levels of IFN-γ secretion when human CAR T cells were co-incubated with their cognate synthetic antigens (**Fig. S4c-d**). Of note, IFN-γ secretion was ~4-fold higher for murine αVHH CAR T cells over αSunTag CAR T cells, likely due to the more stable antigen expression observed on tumor cells (**Fig. 1e**).

**Fig. 2.**
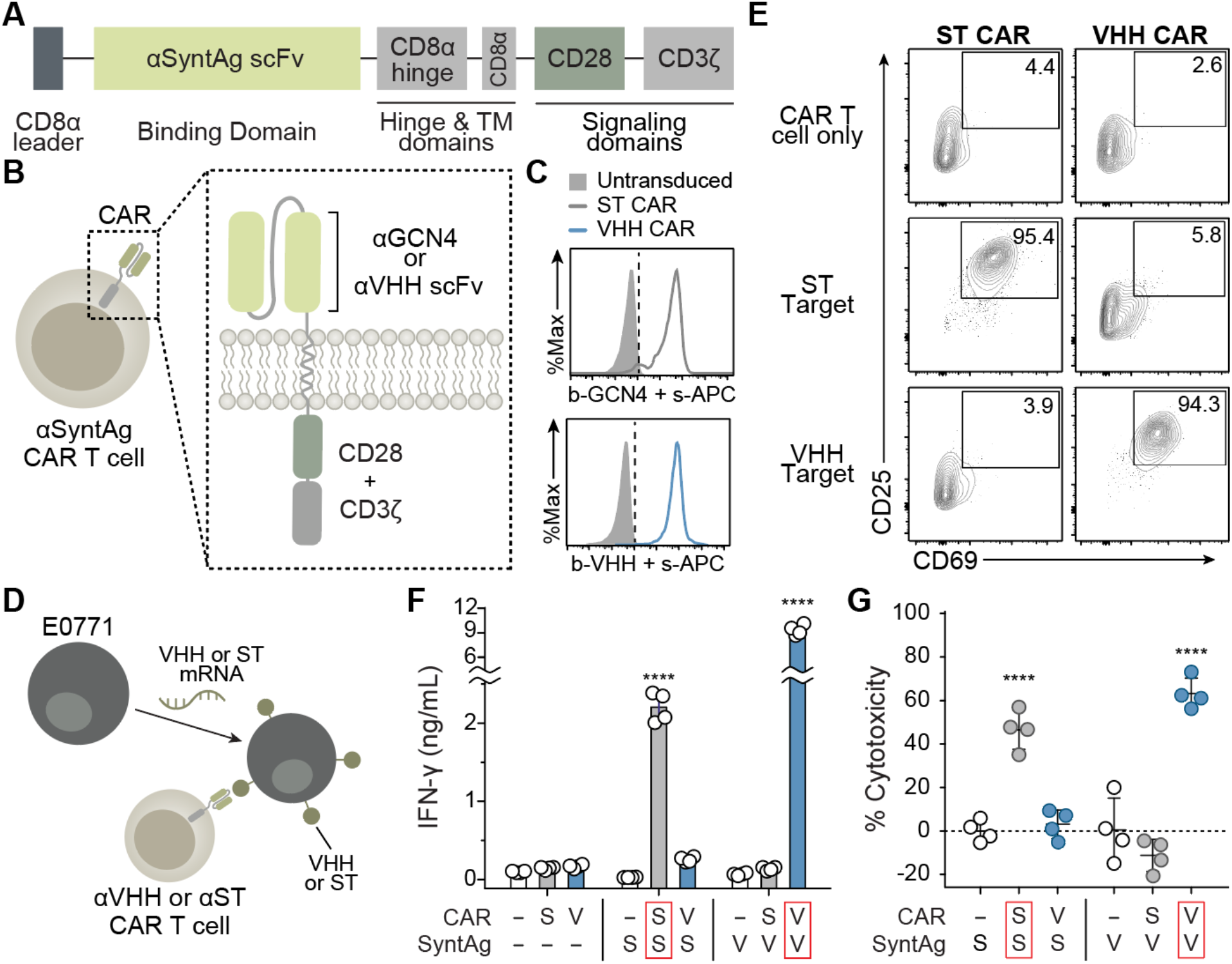
Murine αVHH and αGCN4 CAR T cells recognize and kill tumor cells expressing cognate synthetic antigens. (a-b) Schematic of murine CAR constructs for targeting synthetic antigens expressed on the surface of tumor cells (c) Surface expression of Suntag CAR (top) and VHH CAR (bottom) on primary murine T cells following retroviral transduction. (d) E0771 tumor cells were transfected with VHH or SunTag mRNA and co-incubated with either αVHH or αSunTag CARs. (e) Staining of indicated T cell population with activation markers CD25 and CD69 following co-incubation with ST- or VHH-expressing E0771 tumor cells. (f) Interferon gamma (IFN-γ) secretion by murine αSunTag CAR T, αVHH CAR T or untransduced (WT) T cells following a 24 hr co-culture at a 2:1 effector:target (E:T) ratio with E0771 transfected with either VHH or SunTag mRNA. (g) Killing of transfected E0771 tumor cells following the same 24-hr co-culture. One-way ANOVA; mean ± s.d. is depicted; n = 4; \*\*\*\**p* < 0.0001.

Next, we assessed the ability of both αVHH and αSunTag CARs to kill tumor cells expressing their cognate synthetic antigen. At a 2:1 effector to target ratio, no cytolytic activity was observed by untransduced T cells on either SunTag- or VHH-transfected E0771 tumor cells or by CAR T cells cultured on tumor cells expressing a mismatched synthetic antigen (i.e., αSunTag CAR T cells on VHH-transfected tumor cells, αVHH CAR T cells on SunTag-transfected tumor cells). When αVHH and αSunTag CAR T were co-incubated with their cognate synthetic antigen, however, we observed significant increases in cytotoxicity for both murine (\*\*\*\**p*<0.001, αVHH CAR: 63.2%, αST CAR: 46.6%, **Fig. 2g**) and human CAR T cells (\*\*\*\**p*<0.001, αVHH CAR: 71.3%, αST CAR: 43.7–51%, **Fig. S4e-f**). Both murine and human VHH CAR T cells killed target cells more potently than their SunTag CAR T cell counterparts (**Fig. 2g, Fig. S4e-f**), consistent with earlier data which showed higher secretion of IFN-γ by αVHH CAR T cells (**Fig. 2f, Fig. S4c-d**). Collectively, our data show that tumor cells expressing synthetic antigens can be recognized and killed by CAR T cells. Based on these data, combined with expression stability of VHH, we selected the VHH synthetic antigen and αVHH CAR T cells for subsequent *in vivo* validation.

### αVHH CAR T cells enhance antitumor immunity against solid tumors

A key concern with antigen selection and the development of CAR T cells against them is toxicity in off-tumor tissues (*4*). We postulated that αVHH CAR T cells would exhibit minimal off-tumor toxicity since its target antigen is not natively found in mice. To test this, we adoptively transferred primary murine αVHH CAR T cells intravenously (i.v.) into immunocompetent and naïve C57BL6/J mice and subsequently collected blood serum 7 days post transfer for analysis. Compared to saline controls or mice receiving the equal amounts of untransduced T cells, we found no significant elevations in serum chemistry values, including total protein, aspartate transaminase (AST), alanine aminotransferase (ALT), urea nitrogen (BUN), creatinine, phosphorous, and calcium (**Fig. 3a, Fig. S5**). These data show that αVHH CAR T cells are well tolerated systemically, with no appreciable abnormalities in renal and liver function parameters observed. We further tested whether surface expression of the VHH antigen on its own would affect tumor growth. E0771 and MC38 tumors transduced to stably express VHH do not exhibit altered tumor growth kinetics compared to wildtype tumors (**Fig. 3b, Fig. S6**). Overall, these data show that, independently, both the VHH synthetic antigen and αVHH CAR T cells remain inactive *in vivo*.

**Fig. 3.**
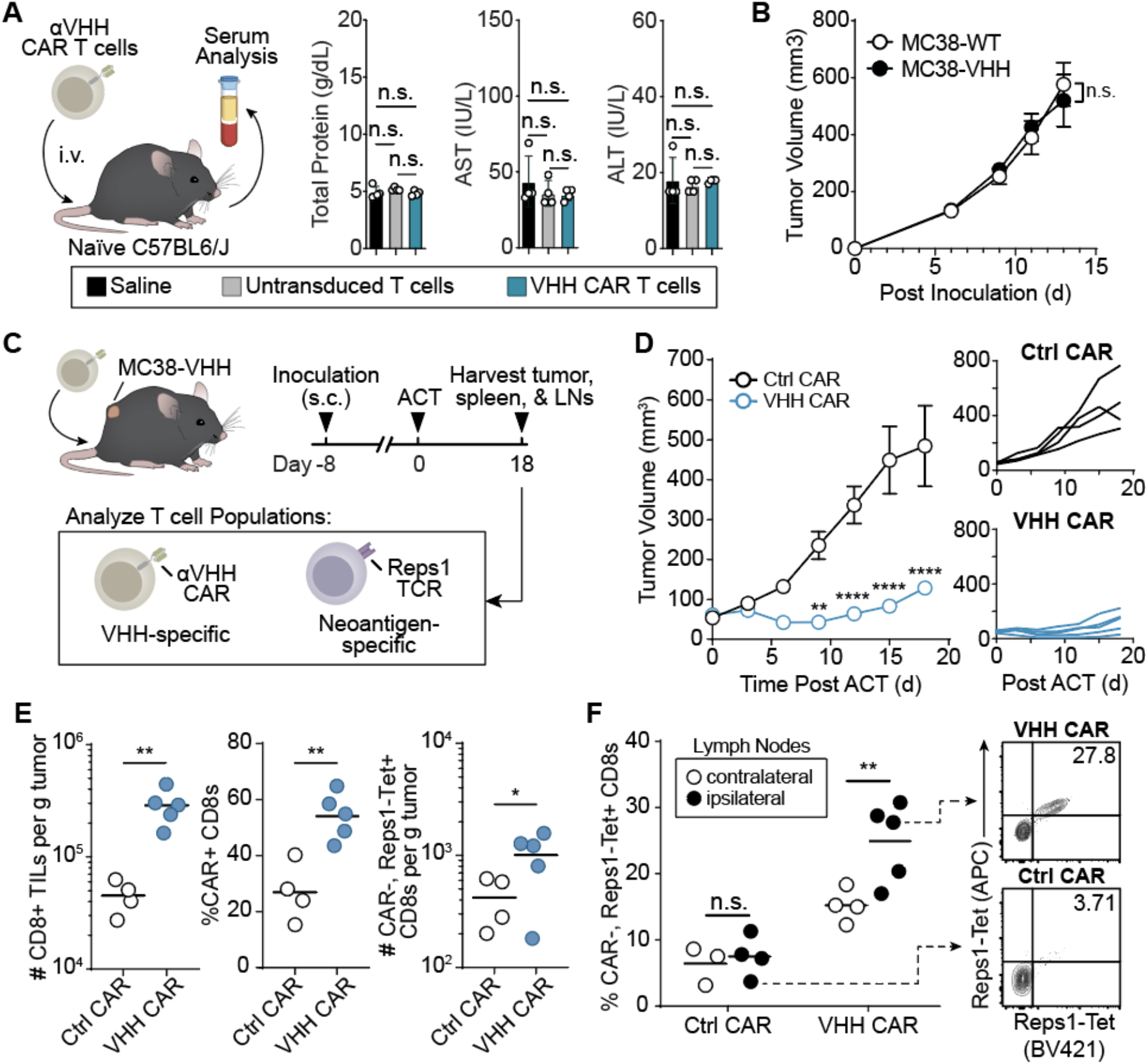
Adoptive transfer of αVHH CAR T cells into mice with VHH-expressing tumors delays tumor growth, promotes infiltration of tumor-reactive T cells in the tumor, and leads to an increased frequency of tumor reactive T cells in the lymph nodes. (a) Blood serum analysis 7d post i.v. administration of αVHH CAR T cells, untransduced T cells, or saline into naïve C57BL6/J mice. One-way ANOVA; mean ± s.d. is depicted; n = 4; n.s. = not significant (b) Tumor growth curves of wildtype MC38 (MC38-WT) or MC38 cells transduced to stably express VHH (MC38-VHH) without adoptive cell transfer of αVHH CAR T cells. Two-way ANOVA, mean ± s.e.m. is depicted; n = 4-6; n.s. = not significant. (c) Mice bearing MC38-VHH tumors were treated with αVHH CAR T cells. (d) Tumor growth curves of MC38-VHH tumor-bearing mice treated with αVHH CAR T cells. Two-way ANOVA, mean ± s.e.m. is depicted; n = 4. (e) Count and frequency of indicated T cell populations isolated from the tumor. Student’s t-test, mean is depicted; n = 4-5. (f) Representative flow plot of endogenous (CAR-) T cell expression of the Reps1 TCR in tdLNs, followed by (g) the frequency of this population in the tumor, non-draining (ndLN) and tumor draining (tdLN) lymph nodes. Student’s t-test, mean is depicted; n = 4-5; \**p* < 0.05; \*\**p* < 0.01; \*\*\**p* < 0.001; \*\*\*\**p* < 0.0001.

To evaluate the antitumor efficacy of αVHH CAR T cells *in vivo*, we adoptively transferred CAR T cells into immunocompetent mice bearing VHH+ MC38 (MC38-VHH) tumors. Compared to treatment with control (i.e., αSunTag) CAR T cells, transfer of αVHH CAR T cells significantly reduced tumor burden (**Fig. 3c**). Within 18 days of adoptive cell transfer (ACT), tumor volumes within the control group grew on average 9-fold, whereas those treated with αVHH CAR T cells grew ~2.5-fold. To further examine the antitumor response, we performed flow cytometry analysis of tumor-reactive T cell populations in the tumor, spleen, and both the ipsilateral (i.e., tumor draining) and contralateral lymph nodes. αVHH CAR T cell treatment augmented the accumulation of CD8+ tumor infiltrating lymphocytes (TILs) compared to treatment with control CAR T cells (\*\**p*<0.01, **Fig. 3e**). Additionally, CAR+ CD8 T cells comprised a higher frequency of the CD8+ TILs within mice receiving αVHH CAR T cells (55% vs. 22%, \*\**p*<0.01, **Fig. 3e**). In contralateral and ipsilateral lymph nodes, however, there were no significant differences in the frequency of CAR+ CD8s among the two treatment groups (**Fig. S7a**), demonstrating that αVHH CAR T cells primarily respond at the tumor site.

Therapies that stimulate immunogenic cell death promote the release and cross-presentation of tumor antigens in the draining lymph nodes and can prime *de novo* T cell responses, analogous to vaccination (*47–49*). By using the tumor as a source of antigens, *in situ* vaccination strategies, such as the intratumoral delivery of IL-12-encoding RNA (*50*) or combination treatment with Flt3L, radiotherapy, and TLR3 agonist (*51*), result in epitope spread that increase response rates to therapy. To determine whether targeting the VHH synthetic antigen could prime against additional tumor antigens, we next quantified the recruitment of endogenous T cells reactive against Reps1, an H-2D^b^-restricted mutant neoepitope characteristic of the MC38 cell line (*52*). In the tumors of αVHH CAR T cell treated mice, we observed a 2.4-fold increase in accumulation of CAR-negative, Reps1-reactive CD8+ T cells in the tumors (\**p*<0.05, **Fig. 3e**), as well as a higher frequency of the same population in the ipsilateral lymph nodes (\*\*\**p*<0.01, **Fig. 3f**). In tumor bearing mice treated with control CAR T cells, however, we found no significant difference in the frequency of Reps1-reactive T cells between the ipsilateral and contralateral lymph nodes (**Fig. 3f**). These data provide evidence that targeting the VHH synthetic antigen with αVHH CAR T cells leads to direct tumor recognition, and that CAR-mediated killing of VHH+ tumors enhances the endogenous response against untargeted neoantigens.

TNBC tumors lack estrogen and progesterone receptor (ER/PR) as well as HER2 expression, making them unresponsive to endocrine and anti-HER2 therapies (*53*). As such, numerous efforts are underway to expand targeted treatment options for patients with TNBC (*54*). We therefore further tested αVHH CAR T cells in mice bearing E0771 tumors, a TNBC originally isolated from a spontaneous mammary tumor in a female C57BL/6 mouse (*55*). Following adoptive transfer of αVHH CAR T cells (**Fig 4a**), all but one mouse treated with αVHH CAR T cells were complete responders for at least 100 days after ACT (CR = 3/4, blue traces, **Fig. 4b**), while all those which received control CARs reached endpoint criteria (tumor volume > 1,000mm^3^) within 30 days of treatment (CR = 0/4, black traces, **Fig. 4b**). To determine whether treatment established immunological memory against the tumor, either naïve mice or complete responders were rechallenged with wildtype (i.e., VHH-negative) E0771 tumor cells in the contralateral mammary fat pad 45 days after initial therapy (**Fig. 4a**). While naïve mice grew tumors that reached endpoint in 28–32 days, cured mice were resistant to rechallenge with VHH-negative E0771 tumor cells (\*\*\*\**p*<0.0001, **Fig. 4c**), demonstrating the development of long-term immunological memory against unknown tumor antigens. Survival of rechallenged mice was significantly extended by at least 70 days over mice treated with control CAR T cells (\*\**p*<0.01, **Fig. 4d**). Collectively, these data demonstrate potent antitumor activity of αVHH CAR T cells in two distinct syngeneic murine tumor models, as well as their potential to elicit systemic antitumor responses analogous to *in situ* vaccination.

**Fig. 4.**
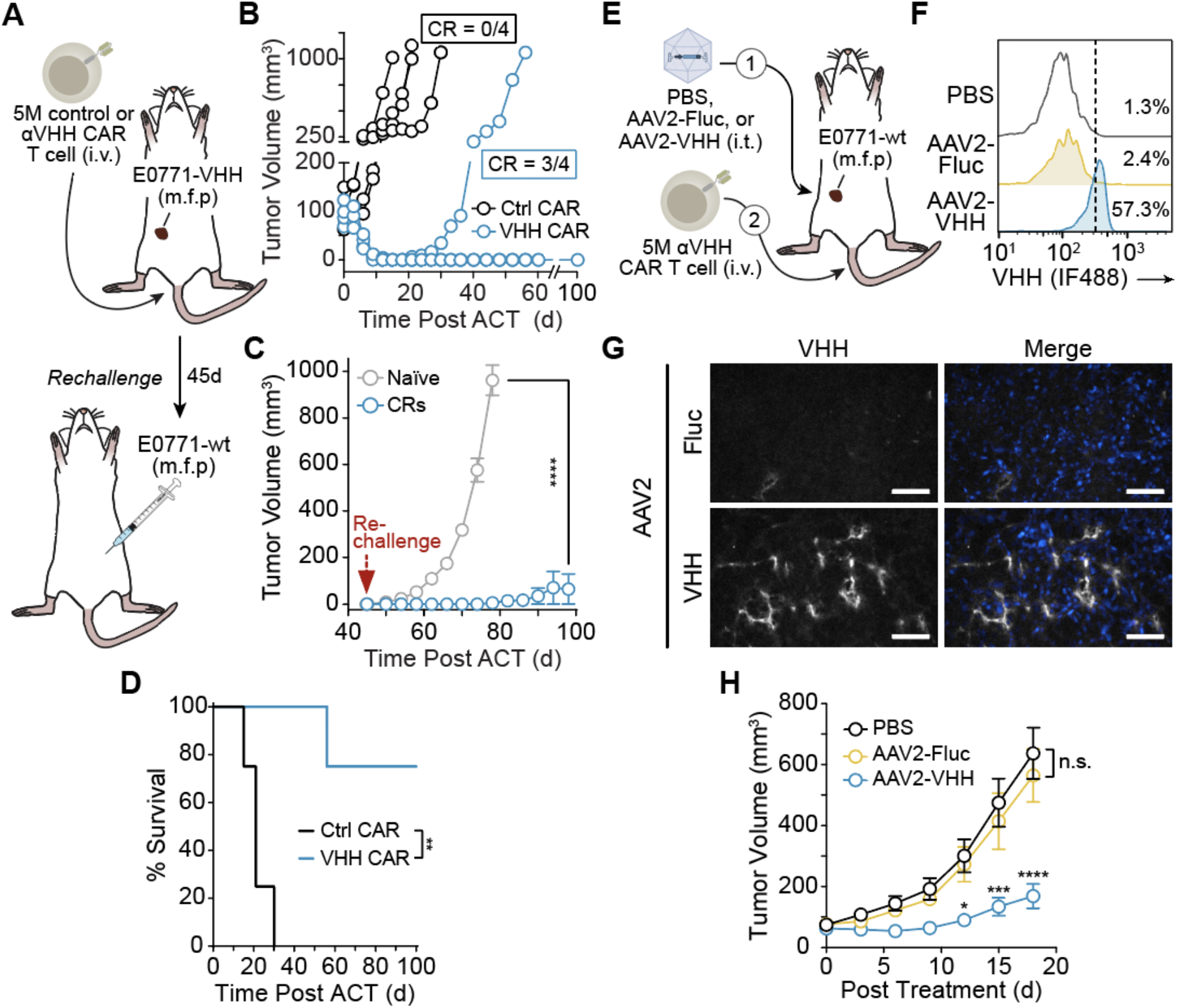
Synthetic antigen treatment promotes potent antitumor immunity in an immunocompetent model of triple negative breast cancer. (a) Mice bearing E0771-VHH tumors were treated with αVHH CAR T cells. (b) Individual traces of tumor growth curves of E0771-VHH tumor-bearing mice treated with control CAR T cells (black) or αVHH CAR T cells (blue) (CR = complete responders). 45 days after initial treatment, cured mice were (c) rechallenged with wildtype E0771 tumor cells. VHH expression on wildtype or transduced MC38 and E0771 tumor cells. Two-way ANOVA, mean ± s.e.m. is depicted; n.s. = not significant. (c) Survival curves of tumor-bearing mice following initial treatment and rechallenge, log-rank (Mantel–Cox) test; \*\**p* < 0.01. (e) Mice bearing wildtype E0771 tumors were first treated with AAV2-Fluc or AAV2-VHH, followed by adoptive transfer of αVHH CAR T cells. (f) Flow cytometry analysis of VHH expression on E0771-Thy1.1 dissociated tumor cells 48 hrs following AAV treatment of 6e9 GCs. Plot is gated on Thy1.1+ cells. (g) Tumor cryosections were fixed and stained for VHH expression. Scale bar = 60 μm. (h) Tumor growth curves of wildtype E0771 tumor-bearing mice treated with AAV2 (Fluc or VHH) and αVHH CAR T cells. Two-way ANOVA, mean ± s.e.m. is depicted; n = 6; \**p* < 0.05; \*\*\**p* < 0.001; \*\*\*\**p* < 0.0001.

### AAV-mediated expression of VHH followed by treatment with αVHH CAR T cells leads to a potent antitumor response

We next sought to explore whether *in situ* delivery of synthetic antigens could sensitize wildtype tumors to CAR T cell cytotoxicity (**Fig. 4e**). Several different viral and nonviral delivery systems are available for effective gene delivery *in vivo* (*56*). Here, we generated an adeno-associated viral (AAV) vector containing GPI-anchored VHH driven by a CMV promoter (AAV2-VHH), as well as a control vector expressing firefly luciferase (AAV2-Fluc), to enable direct delivery of the VHH synthetic antigen to tumors. In MC38 and E0771 tumors expressing a Thy1.1 reporter, AAV2-mediated delivery of VHH led to expression of VHH on the surface of tumor cells as measured flow cytometry staining of Thy1.1+ cells 48 hours after i.t. delivery (E0771 = 57.3% VHH+, MC38 = 37.9% VHH+, **Fig. 4f, Fig. S8**). These results are consistent with immunofluorescence analysis of tumor sections from mice bearing wildtype E0771 (E0771-wt) tumors injected intratumorally with either AAV2-VHH or AAV2-Fluc, further confirming VHH expression in AAV2-VHH treated tumors (**Fig. 4g**). Finally, we combined AAV delivery of the VHH synthetic antigen with systemic administration of αVHH CAR T cells. Established E0771-wt tumors were first treated with an i.t. injection of PBS, AAV2-Fluc, or AAV2-VHH followed by an i.v. injection of αVHH CAR T cells (**Fig. 4e**). Whereas treatment with AAV2-Fluc + αVHH CAR T cells did not significantly affect tumor growth kinetics compared to PBS + αVHH CAR T cell controls, we observed treatment with AAV2-VHH + αVHH CAR T cells led to a significant delay in tumor growth (\*\*\*\**p*<0.0001, **Fig. 4h**). Likewise, we observed a significant reduction in tumor burden for wildtype MC38 tumors only in the case where mice were treated i.t. AAV2-VHH + i.v. αVHH CAR T cells (**Fig. S9**). These data demonstrate that delivery of the VHH synthetic antigen combined with αVHH CAR T cell therapy sensitizes wildtype tumors to immune recognition and significantly improves antitumor responses *in vivo*.

## DISCUSSION

The scarcity of tumor-specific target antigens limits the clinical use of CAR T cells for the treatment of solid tumors. In light of the difficulty in identifying CAR targets for solid tumors, we repurposed the variable-domain-only immunoglobulin fragment VHH as a synthetic antigen. To tether VHH to the outer surface of tumor cells, we designed and tested several genetically encoded constructs consisting of a membrane anchor linked to a CAR recognition domain and demonstrate stable expression of VHH on the surface of various tumor types, including breast, colon, and lung cancer, following mRNA or AAV delivery. Moreover, to provide T cells the ability to recognize VHH-expressing tumors, we designed and validated αVHH CAR T cells, which showed potent antigen-dependent activation, cytokine production, and cytotoxicity upon recognition of VHH on the surface of tumor cells *in vitro*. Importantly, αVHH CAR T cells exhibited potent antitumor activity *in vivo* against VHH+ tumors in syngeneic models of colorectal and TNBC, which led to recruitment of endogenous tumor-reactive cells, resistance to rechallenge, and extended survival.

Many tumor antigen targets identified to date share expression with normal tissue. Consequently, collateral damage associated with low-level expression of TAAs in healthy organs is a safety concern in the use of targeted therapies. Treatment of metastatic colon cancer with αHER2 CAR T cells, for example, led to severe respiratory distress minutes after treatment, likely due to low levels of HER2 expression in the lung, which ultimately developed into to multi-organ failure and death (*21*). Targeting the melanoma TAA, MART-1, with MART-1 specific T cells induced hearing loss and vestibular dysfunction in over half of treated melanoma patients due to cross reactivity with healthy melanocytes found in the inner ear (*57*), while treatment with carboxy-anhydrase-IX (CAIX) targeted CAR T cells led to severe liver toxicity due to shared CAIX expression with epithelial cells lining the bile ducts (*58*). Moreover, multiple neurotoxicity-related patient deaths halted a phase 1 clinical trial of αPSMA CAR T cells for the treatment of metastatic prostate cancer (NCT04227275). The small proportion of healthy neurons expressing PSMA raises the possibility that on-target, off-tumor killing may have contributed to the observed neurotoxicity (*59*). Similarly, single-cell RNA-sequencing data revealed the expression of CD19 in human brain mural cells (*20*), which provides a possible explanation for the high incidence of severe neurotoxicity reported for patients infused with CD19-targeted CAR T cells (*60*).

An alternative to targeting endogenous antigens is to directly deliver exogenous targets to the tumor. This strategy circumvents the need for antigen discovery and enables the use of validated targets to redirect T cell immunity against solid tumors. To this end, several approaches were developed, including antibody-mediated delivery of immunodominant viral epitopes (*61*), intratumoral delivery of sfGFP using tumor colonizing bacteria (*E. coli* Nissle 1917) (*30*), and the local delivery of CD19 using oncolytic virus (OV), for subsequent treatment with their cognate CAR T cells (*29, 31*). Our approach combines AAV-mediated intratumoral delivery of VHH, a xenogeneic protein we engineered for cell surface expression, with VHH-targeting CAR T cells to enable immune recognition. Notably, AAVs are FDA-approved as gene transfer vectors (*62*), while the clinical use of the anti-von Willebrand Factor VHH as a neutralizing agent (*35*) demonstrates that this class of molecules has the potential for safe clinical translation. The data presented in this study demonstrate that αVHH CAR T cells are well tolerated *in vivo*, as measured by serum chemistry in non-tumor bearing mice 7 days after infusion. Nonetheless, preclinical murine models may not fully recapitulate CAR T cell cytotoxicity in humans (*63*) and thus, as with any new CAR T cell construct under consideration, future studies are warranted to assess the potential off-tumor activity of αVHH CAR T against healthy human tissue.

Intratumoral administration of immunotherapies offers an opportunity to maximize therapeutic index by increasing bioavailability and reducing systemic exposure of potent biologics. We envision that intratumoral delivery of VHH could be used to kickstart antitumor immunity, analogous to *in situ* vaccination strategies such as intratumoral delivery of TLR agonists, radiation, and cytokines which use the tumor as its own vaccine to prime or enhance the antitumor immune response (*64–67*). This is supported by our animal studies where we observed that targeting the VHH synthetic antigen enhanced the endogenous response against untargeted neoantigens in a murine model of colon adenocarcinoma, and in a model of TNBC, where we showed that cured mice were resistant to rechallenge with wildtype tumors. Since VHH is a genetically encoded synthetic antigen, it is also amenable for delivery using viral and nonviral gene delivery vehicles beyond AAVs (*56*), including oncolytic viruses and mRNA-loaded lipid nanoparticles (*68, 69*). These modalities offer clinically validated alternatives capable of achieving high delivery efficiency to tumors. Even so, it is unlikely that any single gene delivery vehicle will achieve full tumor coverage, raising the possibility that synthetic antigen-negative variants may escape recognition. Our data revealed, however, that targeting the VHH synthetic antigen with CAR T cells elicits endogenous antitumor responses and promotes long-term immunity against wildtype tumors, indicating that therapeutic responses can be achieved without full tumor coverage.

Together, our studies provide support for the use of VHH as a synthetic antigen to promote CAR-mediated tumor toxicity. Looking forward, our strategy provides an opportunity to target wide array of solid tumors with otherwise limited antigen targets.

## ACKNOWLEDGEMENTS

We thank Dr. D.R. Meyers (Emory) and C.D. Sago for helpful insights.

## Funding

NIH Director’s New Innovator Award DP2HD091793

National Center for Advancing Translational Sciences UL1TR000454

Shurl and Kay Curci Foundation

NIH Shared Instrumentation Grant 1S10OD016264-01A1

National Science Foundation Grant ECCS-1542174

Alfred P. Sloan Foundation (LG)

NIH GT BioMAT Training Grant 5T32EB006343 (LG)

NSF-GRFP Grant No. DGE-1650044 (LG, AS, SND)

Burroughs Wellcome Fund (GAK)

Postdoctoral fellowship from the Wallace H. Coulter Department of Biomedical Engineering and the College of Engineering at Peking University (FS)

## Competing interests

G.A.K. is co-founder and equity shareholder of Glympse Bio and consults for Glympse Bio and Satellite Bio. This study could affect his personal financial status. The terms of this arrangement have been reviewed and approved by Georgia Tech in accordance with its conflict-of-interest policies. L.G., A.Z., D.V., P.J.S. and G.A.K. are listed as inventors on a patent application pertaining to the results of the paper. The patent applicant is the Georgia Tech Research Corporation. The application number is PCT/US21/54972.

