## Supplementary Information for "Sensitizing solid tumors to CAR-mediated cytotoxicity using synthetic antigens"

### Materials and Methods

Animals

Six- to twelve-week-old female mice were used for all experiments. C57BL/6J mice were purchased from the Jackson Laboratory (000664, Bar Harbor, ME, USA) and NSG mice were bred in-house at the Georgia Tech Physical Research Laboratory using breeding pairs purchased from the Jackson Laboratory (005557). All protocols were approved by the Georgia Tech Institutional Animal Care and Use Committee (IACUC). Tumor dimensions were measured with calipers in three dimensions and reported as an ellipsoidal volume.

Cell culture and generation of cell lines

Human embryonic kidney (HEK) 293T cells were obtained from ATCC, the murine E0771 breast cancer cell line was kindly provided by Dr. Susan Thomas (Georgia Institute of Technology), and the murine MC38 colon carcinoma cells were a kind gift from the NCI and Dr. Dario Vignali (University of Pittsburgh). HEK293T and MC38 cells were maintained in Dulbecco's Modified Eagle's Medium (DMEM, Life Technologies 11995073) supplemented with 10% fetal bovine serum (FBS, Thermo Fisher 16140071) and 1% penicillin/streptomycin (Life Technologies, 15140-122). E0771 cells were cultured in RPMI-1640 (Corning 10-040-CV) supplemented with 10% FBS and 1% penicillin/streptomycin. E0771-VHH and MC38-VHH cell lines were generated by lentiviral transduction of wildtype E0771 or MC38 cells, respectively, with the VHH gene driven by the EF1α core promoter in a LeGO-C lentiviral backbone (Addgene #27348). VHH+ cells were single cell sorted using the BD FACS Fusion in the Georgia Institute of Technology's Cellular Analysis Core. Similarly, E0771-Thy1.1 and MC38-Thy1.1 were generated by lentiviral transduction of the Thy1.1 gene, single cell sorted, and stably maintained using Blasticidin (Thermo Fisher A1113903). All cells were cultured at 37°C in 5% $CO_2$.

Construction of αVHH and αSunTag CAR T cells

To obtain the anti-VHH scFv sequence, REmAb® Protein Sequencing with WILD™ analysis (Rapid Novor, Ontario, Canada) was performed on the MonoRab™ Rabbit Anti-Camelid VHH Antibody (Clone 96A3F5, Genscript #A01860). The human CAR is composed of a human CD8α signal peptide, antigen-specific scFv (anti-GCN4[20] or anti-VHH scFv), human CD8α hinge and transmembrane domains, as well as the human 4-1BB and CD3ζ intracellular domains. The murine CAR is composed of a mouse CD8 signal peptide, antigen-specific scFv, mouse CD8α hinge and transmembrane domain, as well as the mouse CD28 and CD3ζ intracellular domains. The CAR constructs were designed to co-express a fluorescent reporter (eGFP) using the T2A sequence and were generated using DNA fragments (custom order from Eurofins Genomics). Human CAR constructs were cloned into a lentiviral vector using the BsiWI and NheI restriction sites. Murine CAR constructs were cloned into a retroviral vector (pMKO.1, kindly provided by Dr. Koichi Araki) using the EcoRI and NotI restriction sites.

*In vitro* transcription of synthetic antigen mRNA

Synthetic mRNA was produced as previously described[74]. Briefly, plasmids containing the template sequence were ordered from Genscript. Plasmid templates were linearized with NotI-HF (New England Biolabs) overnight at 37°C, purified the following day by sodium acetate precipitation and subsequently rehydrated with nuclease-free water. In vitro transcription (IVT) was performed using the HiScribe T7 Kit (NEB) following the manufacturer's instructions, using complete substitution of uridine for N1-methylpseudouridine-5'-triphosphate. The RNA product was treated with DNase for 30 min to remove the template and purified using lithium chloride precipitation. RNAs were capped using guanylyl transferase and 2′-O-methyltransferase (Aldevron) to generate a Cap-1 structure, purified by lithium chloride precipitation, treated with alkaline phosphatase (NEB), and re-purified. The concentration of the purified mRNA was measured using a Nanodrop and subsequently stored at −80°C at stock concentrations of 1-4 mg/mL. Purified RNA product was analyzed by gel electrophoresis to ensure purity.

*In vitro* kinetics of expression

Synthetic antigen encoding mRNA was transfected into target cell lines using jetMESSENGER® mRNA transfection reagent (Polyplus-Transfection, 150-07) following manufacturer's protocol. Following transfection, expression kinetics were analyzed via flow cytometry. Briefly, cells were stained for VHH or SunTag expression and analyzed using the Accuri™ C6 Plus flow cytometer (BD Biosciences) at indicated time points. To isolate extracellular vesicles, target cells were cultured in suitable culture media containing 10% exosome depleted FBS (Gibco, A2720803) prior to transfection. After 24 hours, cell culture supernatant was collected and centrifuged at 2000xg for 30 minutes to pellet cell debris. Cleared supernatant was then transferred to a new tube and 0.5 volumes of Total Exosome Isolation Reagent (ThermoFisher Scientific, 4478359) was added and incubated overnight at 4°C. The next day, extracellular vesicles were pelleted by centrifugation at 10,000xg for 1 hour at 4°C. Extracellular vesicles were then resuspended in PBS before addition to wild-type target cells.

Lenti- and retroviral production

Plasmid DNA was purified using with the E.Z.N.A.® Endo Free Plasmid Maxi Kit (Omega Bio-Tek D6926-03). Recombinant retrovirus was made by co-transfection with pCL-Eco (Imgenex, San Diego, CA) and pMKO.1 retroviral vectors encoding for murine CARs in HEK293T cells using TransIT-293 (MIR2705, Mirus). Virus containing supernatant was collected 48 hrs later, filtered through a 0.45 µm syringe filter (Pall Acrodisc, #4654) to remove cell debris, mixed with Retro-Concentin Virus Precipitation Solution (RV100A-1, System Biosciences, Palo Alto, CA), and stored overnight at 4°C. The next day, retroviral mixture was concentrated at 1500xg for 30 min at 4°C, resuspended in murine T cell media, and immediately used for transduction of primary murine T cells.

Lentivirus was produced by co-transfection of lentiviral expression plasmids with psPAX2 (Addgene #12260) and pMD2.G (Addgene #12259) using TransIT-LT1 transfection reagent (Mirus Bio MIR2300) and HEK293T cells. Viral supernatant collected after 48 hrs, was concentrated using PEG-it™ Virus Precipitation Solution (LV825A-1, System Biosciences, Palo Alto, CA) following the manufacturer's protocol and stored at -80°C until use.

Primary murine CAR T cell production

Cells from Pmel-1 or P14 mouse spleens and lymph nodes were harvested by gently dissociating the tissues using frosted glass slides. Cells were centrifuged at 1000x*g* for 5 min and resuspended in 1x RBC lysis buffer (420301, Biolegend) for 5 min at 4°C. Following the addition of 1x PBS to quench the lysis reaction, cells were centrifuged again, resuspended in complete murine T cell media (cTCM; RPMI + 10% FBS + 1% Pen/Strep + 1x NEAA + 1x Sodium Pyruvate + 50 µM

Beta-mercaptoethanol) containing 100 IU/mL of recombinant human IL-2 (TECIN™ Teceleukin, Bulk Ro 23-6019, National Cancer Institute, Frederick, MD), and passed through a 40 μm cell strainer (732-2757, VWR) prior to counting. Cells were cultured in the presence of either 1 μM human $gp100_{25\text{-}33}$ (Pmel-1) or $gp33_{33\text{-}41}$ (P14). On day 2, cells were collected, washed, and resuspended at $8x10^6$ cells/mL in concentrated retroviral supernatant supplemented with 100 IU/mL hIL2 and 8 μg/mL polybrene. Spinfection was performed in a U-bottom 96-well plate at 2,000x*g* for 90 min at 32°C. Transduced cells were resuspended and maintained at $1x10^6$ cells/mL in fresh cTCM supplemented with 100 IU/mL hIL-2 and passaged daily until use for *in vitro* or *in vivo* experiments on day 6. CAR expression was evaluated by surface staining with biotinylated antigen (biotinylated VHH, Chromotek #gtb-250; biotinylated GCN4, synthesized in-house) followed by a secondary stain with streptavidin-APC (Thermo Fisher #S868). For staining, biotinylated SunTag was synthesized on Rink Amide ProTide (LL) resin using CEM Liberty Blue, including Fmoc deprotection in piperidine, amino acid coupling in N,N'-diisopropylcarbodiimide and Oxyma Pure, N-terminal biotinylation by biotin p-nitrophenyl ester in presence of Oxyma Pure. The peptide was cleaved off resin in trifluoroacetic acid, precipitated in diethyl ether, dried overnight, and resuspended in 10 mg/mL in water for storage at −20°C.

Primary human CAR T cell production

Peripheral blood was drawn from healthy human donors, as approved by the Georgia Tech and Emory University Institutional Review Boards (IRB #H20288). PBMCs were isolated using Lymphoprep density gradient medium (STEMCELL Technologies, 07801) and SepMate-15mL tube (STEMCELL Technologies, 85415), according to manufacturer's intructions. CD3+ cells were then isolated using the EasySep Human CD3 Positive Selection Kit II (STEMCELL Technologies, 17851) and activated using Dynabeads (ThermoFisher, 11131D) at a 3:1 bead-to-cell ratio. Activated cells were cultured in complete human T cell media (hTCM; X-vivo 10 [Lonza #04-380Q], 5% Human AB serum [Valley Biomedical, HP1022] ,10 mM N-acetyl L-Cysteine [Sigma A9165] , 55 uM 2-mercaptoethanol [Sigma, M3148-100ML] supplemented with 50 U/mL recombinant human IL-2 (TECIN™ Teceleukin, Bulk Ro 23-6019, National Cancer Institute, Frederick, MD) for 24 hours at 37°C in 5% $CO_2$. To transduce the activated human T cells, concentrated lentivirus (MOI=25) was added to a 24-well suspension culture plate coated with Retronectin (Takara, T100B) according to the manufacturer's instructions and subsequently centrifuged at 1,200×g for 90 minutes at 37°C. Following centrifugation, activated human T cells in human T cell media supplemented with 100 units/mL of hIL-2 were added to each well and the plate was spun at 1,200×g for 60 minutes at 37°C. Cells were incubated on the virus-coated plate for 24 hours before expansion, and transduction efficiency was evaluated by surface staining with biotinylated antigen as described above. 7 days after activation, T cells were supplemented with Dynabeads at a 1:1 bead to cell ratio. The beads were removed on Day 9. Cells were maintained at a concentration of $7\times10^5$ to $2\times10^6$ cells/mL until Day 10-14 for use in downstream assays.

Liver function analysis

Naïve C57BL6/J mice were administered either PBS, untransduced T cells, or $5\times10^6$ CAR T cells. 7 days following administration, blood was drawn via the jugular vein and collected in serum separator tubes. Samples were sent out for blood chemistry testing at Antech Diagnostics.

AAV production for *in vivo* delivery

The GPI-anchored VHH gene was synthesized as a custom DNA fragment (Eurofins Genomics). To generate AAV expression vectors, the VHH, GFP or Fluc genes were amplified by PCR and placed under the control of a CMV promoter via restriction enzyme cloning using EcoRI and BamHI in the pAAV-CMV expression vector (Takara #6230). Recombinant AAV2 vector was prepared using an AAVpro Helper Free System (Takara #6230). To produce the AAV9 or AAVDJ serotypes, the pRC-mi342 plasmid encoding for the AAV2 *Rep* and *Cap* genes was replaced with either the AAV9 (Genemedi) or AAV-DJ (Cell BioLabs #VPK-420-DJ) rep-cap plasmids. AAV particles were produced by co-transfecting HEK293T cells with the packaging plasmids (pRC and pHelper) and either pAAV-VHH, pAAV-Fluc, or pAAV-GFP using the calcium phosphate transfection method (Takara #631312). Cells were collected 72 h post transfection and AAV particles were extracted and purified using the AAVpro Purification Kit (Takara #6232) following the manufacturer's instructions. Genomic copy number (GC) of AAVs were determined by qPCR using AAVpro Titration Kit Ver. 2 (Takara #6233).

<u>*In vitro* cytotoxicity assay</u>

Tumor (E0771 or A549) cells were transfected in a 96-well plate with 100 ng of either VHH or SunTag mRNA using jetMESSENGER® transfection reagent kit following manufacturer instructions (Polyplus Transfection, 101000005). Transfected target cells were then co-cultured with untransduced (WT) CAR, αVHH CAR or αSunTag CAR T cells at the indicated effector to target cell ratio for 24 hours. Supernatant was collected after incubation and assayed for interferon gamma (IFN-γ) with IFN-γ Mouse ELISA Kit (Invitrogen, KMC4022) or human IFN-γ ELISA kit (ThermoFisher, EHIFNG) according to the manufacturer's protocol. Following the aforementioned 24-hour co-culture, the ability of WT, αVHH CAR and αSunTag CAR T cells to kill transfected E0771 tumor cells was assessed using a lactase dehydrogenase (LDH) release assay according to the manufacturer's instructions (Abcam #ab197004).

<u>Therapy studies</u>

C57BL6/J mice were shaved and inoculated with either $1\times10^6$ MC38-VHH, $5\times10^5$ E0771-VHH, or $5\times10^5$ E0771-wt tumor cells. For E0771 experiments, cells were resuspended in 30 µL PBS (-/-) and implanted i.d. in the left mammary fat pad (fourth). For MC38 experiments, cells were resuspended in 100 µL PBS (-/-) and implanted s.c. into the left flank. Tumor burden, quantified as 0.52 × length × width × depth, was monitored until average tumor volume was approximately $100mm^3$ before initiating treatment. On treatment day, mice were sublethally irradiated with 500 cGy and $5\times10^6$ CAR transduced pmel-1 splenocytes were adoptive transferred via tail vein injections. Recombinant human IL-2 (rhIL-2) was administered intraperitoneally twice daily for 3 days. Mice were classified as complete (CR) or partial responders (PR), or as having progressive disease (PD) or stable disease (SD) based on the RECIST criteria[75].[75]. On Day 18 post ACT, MC38-VHH tumors and lymph nodes were isolated for flow cytometry analysis. For E0771-VHH experiments, CRs were rechallenged 45 days after initial treatment with $5\times10^5$ synthetic antigen negative (E0771-wt) tumor cells in the right mammary fat pat (fourth).

<u>Flow cytometry analysis of T cells *in vivo*</u>

All antibodies for flow cytometry were purchased from Biolegend. Single cell suspensions from spleens and lymph nodes were prepared by homogenizing the tissue between the frosted end of glass slides. Homogenized cells were passed through a 40 µm cell strainer, depleted of red blood cells using 1x RBC lysis buffer (Biolegend 420302). For tumors, less than 1g of MC38-VHH tumors were enzymatically and mechanically dissociated using the Mouse Tumor Dissociation Kit (Miltenyi, 130-096-730) and gentleMACS Dissociator (Miltenyi, 130-093-235). TILs were then isolated from the single cell suspension using a density gradient with Percoll Centrifugation Media (VWR, 17-5445-01) and DMEM Media (10% FBS, 1% Pen-strep) at a 44:56 volume ratio. The panel used for staining is as follows: anti-CD8-BV786 (53-6.7), anti-Thy1.1-BV711 (IM-7). All antibodies were used for staining at 1:100 dilution from stock concentrations. Cell viability was assessed by staining with LIVE/DEAD™ Fixable Aqua Dead Cell Stain Kit (ThermoFisher Scientific, L34957) following manufacturer's instructions. To detect VHH CAR expression, biotinylated VHH (Chromotek, gtb-250) and streptavidin-Alexa Fluor™ 488 (Invitrogen, S32354) were used for staining at a 1:100 dilution from stock concentrations. Reps1 reactive T cells were detected by Reps1 pMHC tetramer staining. To ensure signal specificity, samples were stained with Reps1 pMHC tetramers separately conjugated to streptavidin-APC (Invitrogen, S868) and streptavidin-BV421 (Biolegend, 405225) following the procedure below. For surface staining, cells were blocked with anti-Fc receptor anti-CD16/CD32, and then stained in FACS Buffer (1x DPBS, 2% FBS, 1 mM EDTA, 25 mM HEPES). Fixation was performed using eBioscience Intracellular Fixation & Permeabilization Buffer Set following the manufacturer's instructions (Thermo, 88-8823-88). Counting beads (Thermo, C36950) were added to each sample of stained cells prior to analysis by the LSR Fortessa Flow Cytometer (BD).

Tetramer production

For tetramer staining, tetramers were generated from pMHC monomers and streptavidin as previously described[76]. Briefly, biotinylated Reps1 pMHC monomers were diluted in exchange buffer (20 mM Tris-HCl, 150 mM NaCl, pH 7.0) at 0.15 mg/mL. APC or BV421 conjugated streptavidin was added to the diluted pMHC monomers in 4 small aliquots to achieve a final 4:1 pMHC to streptavidin molar ratio. Each aliquot was added to the diluted pMHC monomers every 5 min for a total of 4 times. After each addition, the pMHC-streptavidin mixture was mixed thoroughly and incubated on ice for 5 min. 15 min after the last addition, the pMHC tetramer was centrifuged at 16,000xg for 1 min, and the supernatant was collected for cell staining at 1:50 dilution (0.2 μg/sample).

AAV therapy study

C57BL6/J mice were shaved and inoculated with $5x10^5$ E0771-wt tumor cells resuspended in 30 μL PBS (-/-) and i.d. in the left mammary fat pad (fourth). On treatment day (tumor volume ~ $100mm^3$), mice were sublethally irradiated with 500 cGy and injected intratumorally with $6\times10^9$ GCs of either AAV2- Fluc, AAV2-VHH or PBS. 6 hours later, $5x10^6$ αVHH CAR T cells were administered intravenously. Recombinant human IL-2 (rhIL-2) was administered intraperitoneally twice daily for 3 days. Tumor volumes were measured every 3-4 days. For flow analysis, tumors were harvested from mice bearing E0771-Thy1.1 tumors, dissociated and stained as described above using the following panel: anti-Thy1.1-PE (Biolegend, OX-7), anti-VHH-iFluor 488 (Genscript, 96A3F5), LiveDead (info). Analysis was performed by the LSR Fortessa Flow Cytometer (BD). For histological analysis, a subset of tumors were harvested 48 hrs following injection of either PBS, AAV2-Fluc or AAV2-VHH injections. Tumors were frozen in Clear Frozen Section Compound (VWR #95057-838) and were later cryosectioned (8 μm, Emory Histology Facility). Frozen tissues were equilibrated at room temperature for 30 minutes and then incubated in chilled acetone at -20°C. Slides were soaked in chilled 1x PBST (PBS+ 0.1% Tween20) for 5 minutes, transferred to warmed Antigen Retrieval Solution (10mM sodium citrate, pH 6.0) and heated at 95°C for 20 minutes. The slides were then removed and allowed to cool for 20 minutes before being washed with chilled 1x PBST for 10 minutes. A PAP pen was used to draw a hydrophobic barrier around the tissue and the sample was incubated in PBS, 1% bovine serum albumin (BSA, Sigma #A7030-50G) and 10% donkey serum (Jackson Laboratories #017000121) for 60 minutes at room temperature. The solution was decanted, and the tissue was incubated with the primary rabbit anti-VHH antibody (Genscript, 96A3F5) in PBS, 1% BSA and 10% donkey serum in a humidified chamber overnight at 4°C. Following the overnight incubation, the tissue was washed 3 times with chilled PBST, and the sample was incubated with the diluted Donkey anti-rabbit Alexa Fluor® 647 secondary antibody (Jackson ImmunoResearch, 711-605-152) in 1% BSA and 10% donkey serum at room temperature for 60 minutes in the dark. The tissue was then washed three times with chilled PBST for 5 minutes, and the sample was incubated with 1μg/mL of Hoechst stain in PBST for 3 minutes. After incubation, the sample was washed three times with chilled PBS for 5 minutes and the coverslip was mounted using one drop of ProLong Glass Antifade Mountant (Invitrogen #P36982). The coverslip was then sealed and stored in the dark at 4°C until imaged. Images were acquired using a Hamamatsu Flash 4.0 v2 sCMOS camera on a PerkinElmer UltraView spinning disk confocal microscope mounted to a Zeiss Axiovert 200 M body. Images were taken using a 20x NA 0.8 plan-apochromat objective lens (Zeiss). Z-stack images were acquired using Volocity (PerkinElmer) with an ASI PZ-2150 motorized piezoelectric stage with 0.5 μm increments. Linear contrast enhancement was applied identically to all images for clarity.

Statistical analysis

Appropriate statistical analyses were performed using GraphPad Prism (*$P < 0.05$, ** $P < 0.01$, *** $P < 0.001$, **** $P < 0.0001$). Central values represent mean and error bars depict s.e.m. Flow cytometry data were analyzed using FlowJo X (FlowJo, LLC). Power analyses were performed using G*Power 3.1 (HHUD).

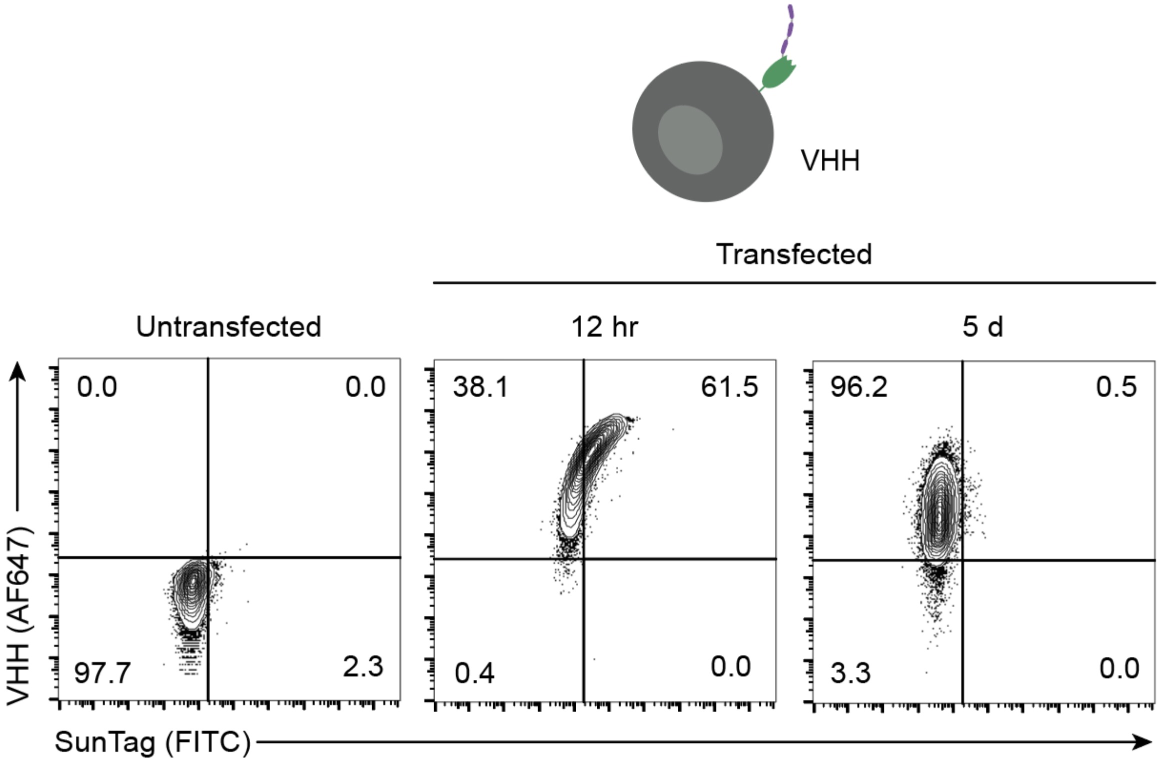


**Fig. S1.** Flow cytometry staining for VHH and SunTag in A549 cells following transfection of GPI-anchored, VHH-linked SunTag. Analysis was performed 12 hours and 5 days post transfection.

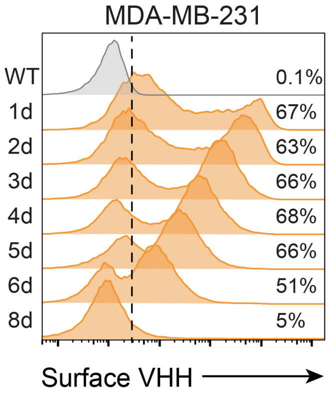


**Fig. S2.**Timecourse of expression of GPI-anchored VHH on the surface of MDA-MB-231 cells following mRNA transfection.

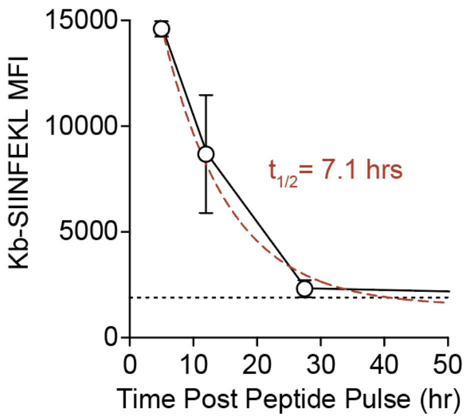


**Fig. S3.** Surface expression kinetics of the Kb-SIINFEKL pMHC complex on the surface of MC38 tumor cells following peptide pulsing with the SIINFEKL peptide for one hour at room temperature.

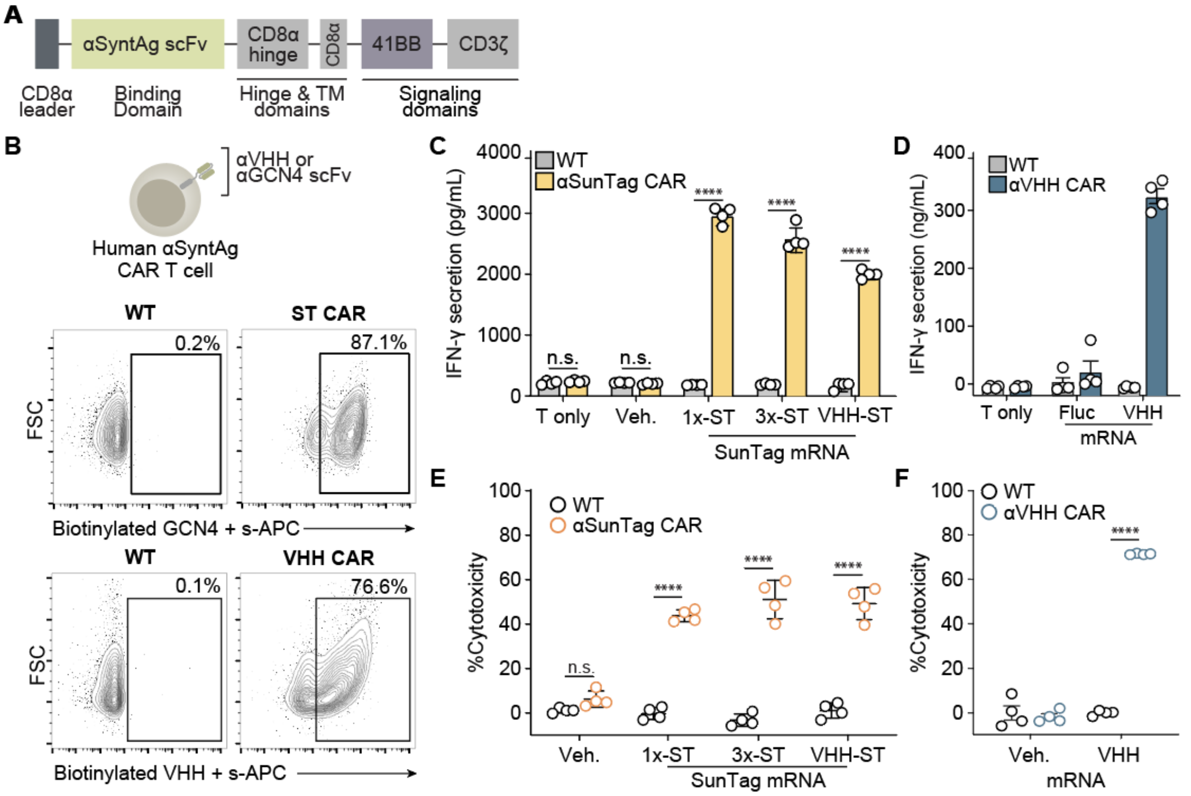


**Fig. S4. Human αVHH and αGCN4 CAR T cells recognize and kill tumor cells expressing cognate synthetic antigens**. (**a**) Schematic of human CAR constructs for targeting synthetic antigens (SyntAg) expressed on the surface of tumor cells. (**b**) Surface expression of SunTag CAR (top) and VHH CAR (bottom) on primary human T cells following lentiviral transduction. (**c**) Interferon gamma (IFN-γ) secretion by αSunTag CAR T or untransduced (WT) T cells following a 24 hr coculture with A549 tumor cells expressing indicated SunTag constructs at a 1:1 effector:target (E:T) ratio (**d**) Interferon gamma (IFN-γ) secretion by αVHH CAR T or untransduced (WT) T cells following a 24 hr coculture with A549 tumor cells expressing VHH at a 2:1 effector:target (E:T) ratio. Killing of A549 tumor cells expressing either (**e**) SunTag constructs or (**f**) VHH following a 24 hr coculture with either CAR or untransduced (WT) T cells. Student's t-test; mean ± s.d. is depicted; n = 4; ****$p < 0.0001$.

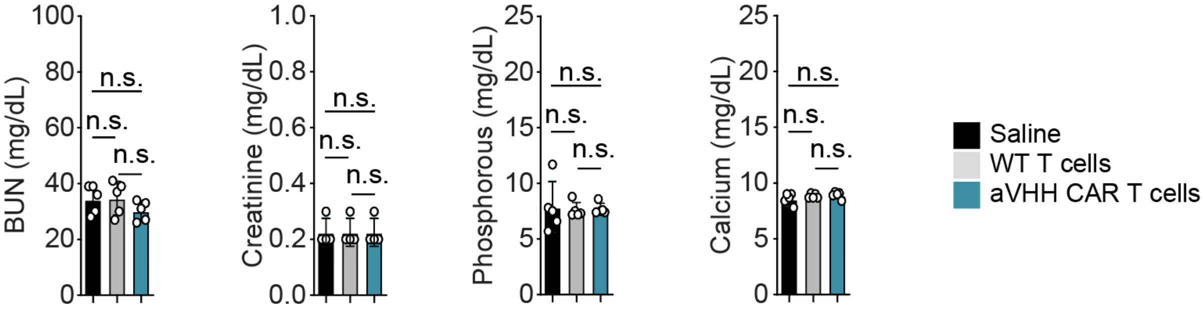


**Fig. S5. Additional Blood Serum Analysis**. (**a**) Additional blood serum analysis 7d post i.v. administration of αVHH CAR T cells, wild-type T cells, or saline.

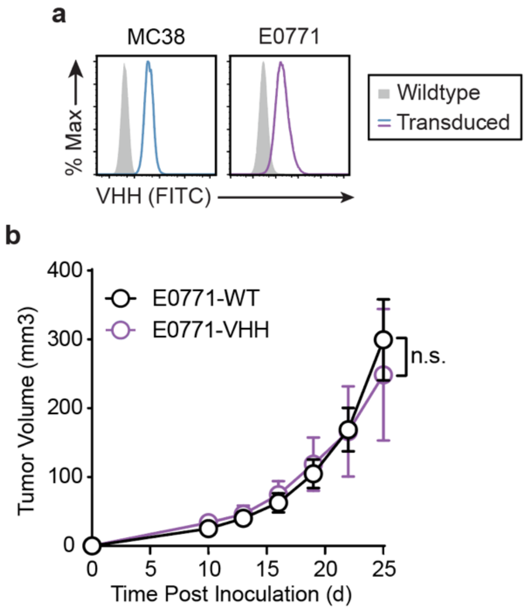


**Fig. S6. Tumor model characterization** (a) VHH expression on wildtype or transduced MC38 and E0771 tumor cells. (b) Tumor growth curves of wildtype E0771 (E0771-WT) or E0771 cells transduced to stably express VHH (E0771-VHH) without adoptive cell transfer of αVHH CAR T cells. Two-way ANOVA, mean ± s.e.m. is depicted; n = 4-6; n.s. = not significant.

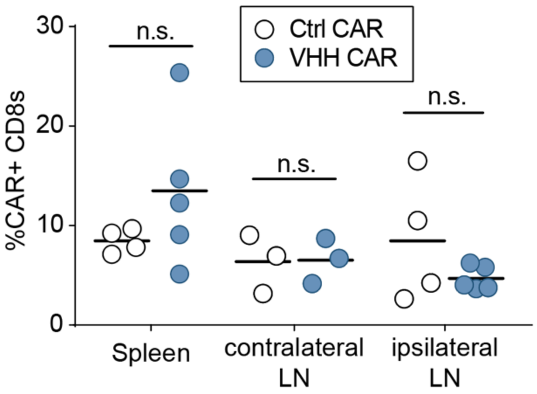


**Fig. S7. Immune cell characterization in lymph nodes of treated mice.** Frequency of CAR+ CD8 T cells isolated from contralateral and ipsalateral LNs in treated mice. (Student's t-test ANOVA, mean is depicted; n = 4-5; n.s. = not significant.

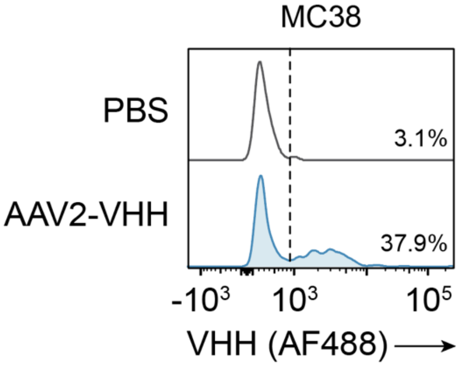


**Fig. S8. AAV-mediated expression of VHH in MC38 tumors.** VHH expression detected by flow cytometry 44 hrs post injection of 1.5e9 GCs of AAV2-VHH into MC38-Thy1.1 tumor cells. Data are gated on Thy1.1 (tumor) cells.

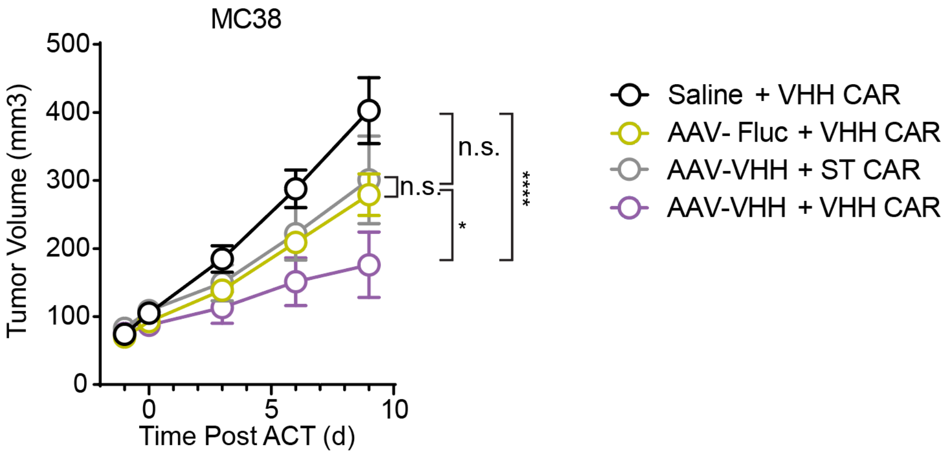


**Fig. S9. AAV-mediated expression of VHH in combination with adoptive transfer of αVHH CAR T cells delays tumor growth in MC38 tumors.** Tumor growth curves of wildtype MC38 tumor-bearing mice treated with AAV2 (Fluc or VHH) and indicated CAR T cells. Two-way ANOVA, mean ± s.e.m. is depicted; n = 6; *$p < 0.05$; ****$p < 0.0001$.
